# Dynamical transitions of the actomyosin cortex can trigger single cell morphogenesis

**DOI:** 10.1101/2021.06.25.450008

**Authors:** Hongkang Zhu, Roberto Alonso-Matilla, Zachary McDargh, Ben O’Shaughnessy

**Author notes:** These authors contributed equally (Hongkang Zhu, Roberto Alonso-Matilla). Correspondence (B.O.).

## Abstract

Morphogenetic changes driven by actomyosin contractile forces are well-characterized at the tissue level. At the single cell level, shape changes steered by actomyosin contractile forces include mitotic rounding and cytokinetic furrow ingression. In some cases, more complex shape transitions associated with spatial patterning of the cortex were observed. The actomyosin cortex was widely studied using active gel frameworks, and stabilized contractile instabilities were shown to generate patterns, but whether complex shapes can emerge from these cortical patterns is not established. Here we show that complex morphogenetic changes at the single cell level can accompany cortical patterns, using a minimal active gel model. For sufficiently low membrane-cortex drag, an initially homogeneous cortex spontaneously develops stripes associated with stable furrows, similar to furrowing observed in cells. Our work suggests that controlled cortical instability can trigger morphogenesis at the cellular level.

## Introduction

Spatiotemporal regulation of actomyosin material densities is used to drive shape changes at multicellular level during tissue morphogenesis in development. For example, planar polarized actomyosin patterns on the apical cell surfaces drive the convergent extension in *Drosophila* germband extension (Rauzi et al., 2010) and chicken neurulation (Nishimura et al., 2012). And tissue-wide graded actomyosin patterns at the ventral side of *Drosophila* embryos are responsible for ventral furrow formation during gastrulation (Heer et al., 2017).

At the single and subcellular levels, spatiotemporal regulation of actomyosin densities is also used to generate cortical flows and shape change. During cell division, the mitotic cell rounding is driven by the retraction and rigidification of the actomyosin cortex (Maddox and Burridge, 2003), following by the cleavage furrow formation by the actomyosin contractile ring (Pollard and O’Shaughnessy, 2019; Turlier et al., 2014). Besides, cell polarity in early *C. elegans* is established from cortical flows that are propelled by gradients in actomyosin contractility (Munro et al., 2004).

More complex patterning of actomyosin densities associated with shape patterning has been observed at the single cell level. For example, parallel stripe patterns of actin modulation simultaneously on the single and multicellular levels were seen in *C. elegans* embryo (Priess and Hirsh, 1986) and *Drosophila* trachea (Hannezo et al., 2015).

In some these cases, the spatial patterning was associated with shape patterning at the single and supracellular levels, with the cell surfaces furrowing in space in *C. elegans* embryo (Priess and Hirsh, 1986) and *Drosophila* trachea with *kkv* mutation (Hannezo et al., 2015). A complex shape transition associated with actomyosin patterning has also been seen at the subcellular organelle level. In *Drosophila* salivary glands, myosin stripes were seen on the actin-coat secretory granules, which are closely correlated with the pronounced furrows on the granule membrane (Rousso et al., 2016).

How shape variation emerges in these systems at the single cell level has not been quantitatively explained. The actomyosin cortex of the cell has been widely described by active gel models (Berthoumieux et al., 2014; Mietke et al., 2019; Salbreux and Julicher, 2017; Turlier et al., 2014). Patterning can occur due to actomyosin contractile instability stabilized. This contractile instability, protected by turnover (Thiyagarajan et al., 2021), can lead to inhomogeneous densities of actomyosin material, because regions with higher density can be self-reinforced by the flow generated by the higher density itself (Bois et al., 2011a). This has been used to explain the experimentally observed patterning in *Drosophila* trachea (Hannezo et al., 2015), Caco-2 epithelial cells (Moore et al., 2014b) and early *C. elegans* embryo (Nishikawa et al., 2017). However, whether the actomyosin contractile instability as described by active gel models can lead to shape changes has not been demonstrated. One theoretical study by Mietke et al. quantitatively described the interactions between active material and shape change, but they had to invoke artificially large bending moduli in order to stabilize the deformed shapes (Mietke et al., 2019).

Here we show that complex morphogenetic changes in the actomyosin cortex are indeed predicted by the active gel framework, associated with patterning changes. We introduce a standard minimal active gel model of the actomyosin cortex. First, we study this model on a plane, and demonstrate using linear stability analysis and numerical simulations that cortical contractility drives spontaneous formation of striped patterns. Then we apply the model on an initially cylindrical surface, and find that the cortex spontaneously assembles into circumferentially oriented stripes, which locally constrict the cortical surface, creating stable furrows. These furrows colocalizing with actomyosin stripes are reminiscent of the deep furrows observed on the exocytotic vesicles in *Drosophila* salivary gland cells (Rousso et al., 2016) and the *Drosophila* tracheal tubes with *kkv* mutation (Hannezo et al., 2015).

## Results

### Active gel model of the cellular actomyosin cortex

Here, we introduce a simple coarse-grained model considering the cortex as an isotropic active gel embedded in an infinitesimally thin layer. The cortex is described at time *t* by a surface **X**(*x*^1^, *x*^2^, *t*), where *x*^1^ and *x*^2^ are the surface coordinates. Following previous works (Berthoumieux et al., 2014; Mietke et al., 2019; Salbreux and Julicher, 2017; Turlier et al., 2014), we introduce an active density field *ρ*(*x*^1^, *x*^2^, *t*) that describes density of cortical actomyosin material at coordinates (*x*^1^, *x*^2^) at time *t*. The mass conservation equation for the density field reads

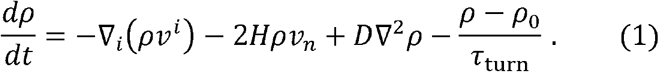

The terms on the right-hand side describe, respectively, advection, aggregation/dilution due to local contraction/expansion of the cortical surface, diffusion, and turnover (Fig. 1A).

**Fig. 1.**
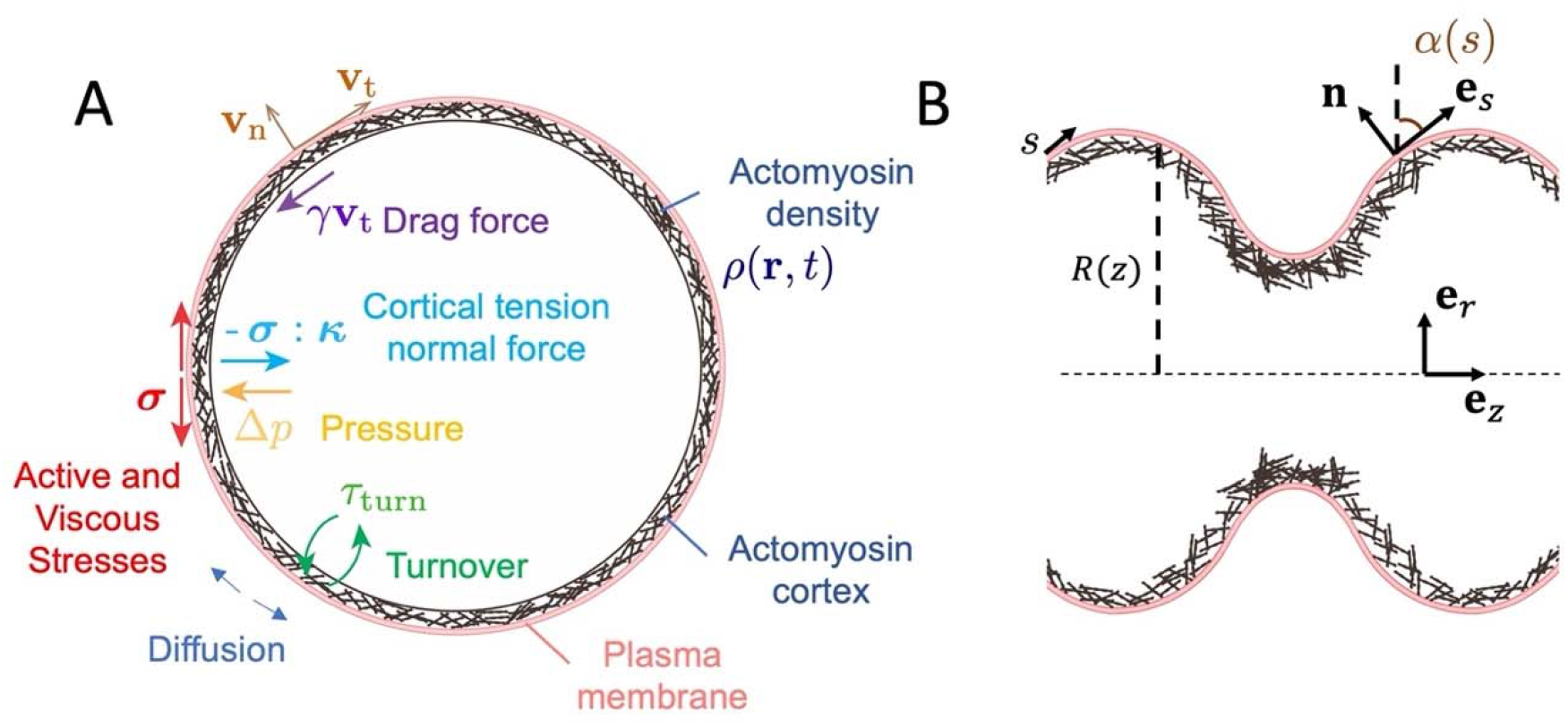
(A) Schematic diagram illustrating our model. The cortex is modelled as a fluid actomyosin surface whose dynamics are determined by force balance. The model includes forces arising from active and viscous stresses, pressure, and drag, as well as the dynamic effects of diffusion, and turnover. (B) Representation of the axisymmetric tubular surface with arc-length parameterization *s*.

The in-plane advection is driven by cortical flows, which emerge from contractile stresses generated by myosin motors as they bind and pull on the actin filament network. Here, ∇_*i*_ denotes the covariant derivative, and *v^i^* is the component of the flow velocity field tangent to the two-dimensional cortex.

The second term on the right-hand side describes changes in density due to local expansion/compression of the cortical surface, where *H* is the local mean curvature of the cortex and *v_n_* is the velocity of the cortex normal to its surface. Local expansion of the cortex (i.e. *Hv_n_* > 0) decreases the cortical density whereas local compression (i.e. *Hv_n_* < 0) increases the cortical density. Note that *H* = (*κ*_1_ + *κ*_2_)/2, where κ_1_ and κ_2_ are the principal curvatures of the surface of the cortex, i.e. the eigenvalues of the curvature tensor *κ_ij_*. The curvature tensor is given by *κ_ij_* = –**n** · *∂_j_***e**_*i*_, where *∂_j_* = *∂*/*∂x^j^* are the partial derivative with respect to the surface coordinates, **e**_*i*_ = *∂_i_***X** are the basis vectors tangent to the surface, and **n** = (**e**_*i*_ × **e**_*j*_)/|**e**_*i*_ × **e**_*j*_| is the unit normal vector of the surface (Fig. 1A–B).

We include the diffusion of actomyosin material. The operator ∇^2^ is the Laplace-Beltrami operator on the cortical surface. The fourth term describes the turnover processes of cortical constituents, where *τ*_turn_ is the turnover time required for the density of a perturbed cortex to relax to the homogeneous steady-state density *ρ*_0_, which depends on association rate *k_a_* and dissociation rate *k_d_* with the relations *ρ*_0_ = *k_a_*/*k_d_* and *τ*_turn_ = 1/*k_d_*.

The normal velocity *v_n_* also determines the dynamics of shape change,

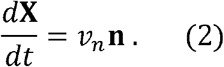

Thus, as active stresses generated by the cortex create out-of-plane forces, the shape of the cortical surface will evolve.

Now, we proceed to analyze the force balance of the cortex. Mechanical stresses are generated by myosin forces, and gradient in stress leads to cortical flows and cellular deformations (Fig. 1A). We take the thin-limit approximation and neglect normal stresses and cytoplasmic viscous dissipation. Tangential force balance on the active gel reads

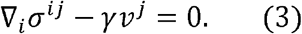

The first term in Eq. (3) describes the effect of internal forces in the gel, and the second term describes cortex-membrane drag forces with drag coefficient *γ*.

For the stress, we consider the actomyosin cortex as a viscous fluid based on the assumption that the time scales of events in the cortex are long enough that only viscous stress is present. We use the following standard constitutive relation for *σ^ij^*

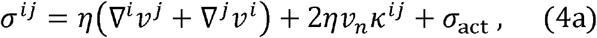

where *σ*_act_ is the active stress assumed to be a function of the cortical density whose value is saturated to the maximum active stress 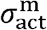 at high *ρ* with a half-saturation density *ρ_s_*,

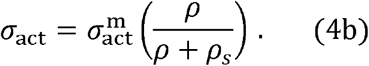

The first term in Eq. (4a) is the gel viscous stress, where *η* is the two-dimensional viscosity of the cortex. Note that, for simplicity, we have assumed that the shear viscosity *η_s_* and the bulk viscosity of the gel *η_b_* are equal *η* = *η_s_* = *η_b_*. The second term in Eq. (4a) describes the viscous contribution to the stress that arises as the curved two-dimensional cortex expands/contracts locally.

Because normal stress is neglected, the force balance normal to the surface reduces to the generalized 3-D Young-Laplace equation

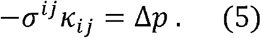

Eq. (5) is the shape equation of the gel and describes the balance between viscous forces, active forces and pressure forces that act normal to the two-dimensional surface of the gel. Δ*p* is the pressure difference between the inside and the outside of the cell. Previous works suggest that the cytoplasm behaves as a poroelastic material, where the time scale for pressure equilibration is on the order of tens of seconds (Charras et al., 2005; Moeendarbary et al., 2013). At longer times compared to this equilibration time, pressure inside the cell can be considered homogeneous. Thus, we assume a uniform cytoplasmic pressure Δ*p*. In our model, we assume constant cellular volume, and Δ*p* plays the role of a Lagrange multiplier so as to impose conservation of the total cell volume.

The parameters we used are summarized in Table 1. The 2D viscosity *η* is estimated *η* = *η*_3D_*h* ≈ 10 pN · s/μm^2^ based on the 3D viscosity *η*_3D_ ≈ 100 pN · s/μm^3^ (Evans and Yeung, 1989) and the thickness of the cortex *h* ≈ 0.1 μm (Chugh and Paluch, 2018). The maximum active stress is assumed to be 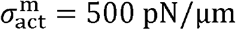 based on cortical tension measurement in fibroblasts (Tinevez et al., 2009). Both the cortex-membrane drag coefficient *γ* and the actomyosin diffusivity *D* are scanned within a range of values.

**Table 1.** Parameters used in simulations

| Parameter | Symbol | Value |
| --- | --- | --- |
| $\eta$ | 2D cortex viscosity | 10 pN · s/μm |
| $\tau_{\text{turn}}$ | Turnover time | 10 s |
| $\sigma_{\text{act}}^{\text{m}}$ | Maximum active stress | 500 pN/μm |
| $\rho_0$ | Homogeneous steady-state density* | $\rho_s/2$ |
| $\gamma$ | Drag coefficient | Scanned |
| $D$ | Diffusivity | Scanned |

\* All densities are scaled by the half-saturation density $\rho_s$

### Conditions for spontaneous pattern formation in the actomyosin cortex

We first apply the model to a cortex anchored to a flat plasma membrane. In this case, both the curvature terms *H* and vanish, and the normal force balance Eq. (5) becomes trivial with *κ_ij_* = 0 and Δ*p* = 0. The normal component of the velocity *v*_n_ is also zero, so there is no shape change.

Consider a uniform actomyosin cortex with the uniform density *ρ*_0_ within a square region of length *L* and periodic boundary conditions. The homogeneous state *ρ* = *ρ*_0_ is a steady-state solution since it satisfies the force balance with zero velocity and zero time derivative. To investigate whether spontaneous pattern formation occurs in this system, we perturb the density *ρ* = *ρ*_0_ + *δρ* exp(*i***k** · **x** + *λt*) with a small amplitude *δρ* ≪ *ρ*_0_, where **k** is the wave vector and *λ* is the growth rate of the perturbation. We assume the perturbed velocity has the form **v** = **δv** exp(*i***k** · **x** + *λt*). The linear stability analysis presented below will reproduce results of Moore et al. and Hannezo et al. (Hannezo et al., 2015; Moore et al., 2014a)

Now we write the active stress of Eq. (4b) as 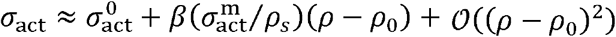 to first order in *ρ* – *ρ*_0_, where 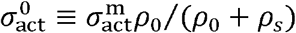 and 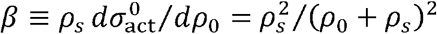 is the dimensionless susceptibility of active stress with respect to density in the homogeneous state (see Eq. (4b)). Using these above forms for *ρ* and **v** in Eq. (4a) to obtain the stress, *σ^ij^*, the force balance Eq. (3) yields

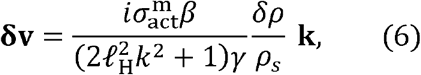

where 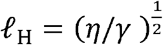 is the hydrodynamic length. Below the hydrodynamic length *ℓ*_h_, viscous dissipation of active stress dominates; above this scale, frictional dissipation dominates. Using this relation, we determine the growth rate *λ* from the time evolution of density, Eq. (1). Retaining first order terms in *δρ* and **δv** yields the dispersion relation

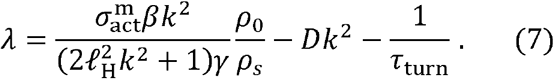

A positive *λ* suggests pattern formation with wavenumber *k*. The relation in Eq. (7) suggests that lower cortex-membrane drag coefficient *γ* and lower actomyosin diffusivity *D* destabilize homogenous states and favor spontaneous pattern formation. To find if there is such a wavenumber with positive growth rate, we maximize *λ* with respect to *k* and we find the most unstable wavenumber *k*_max_ obeys the relation

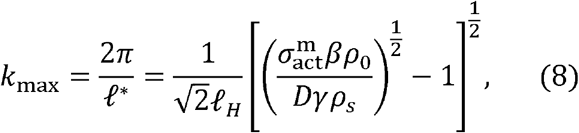

with the maximal growth rate

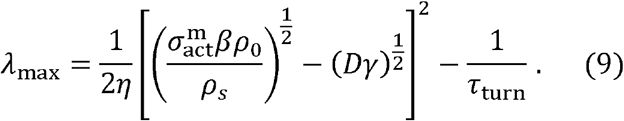

The lengthscale corresponding to *k*_max_ in the predicted characteristic length scale of formed patterns, *ℓ*\* of Eq. (8). This reproduces the estimate of the pattern length scale from Hannezo et al. (Hannezo et al., 2015).

We now study the conditions necessary for pattern formation. The requirement is that the highest growth rate *λ* be positive. From Eq. (9),

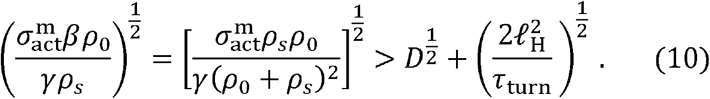

This inequality defines a phase diagram expressing which values of drag coefficient *γ* and diffusivity *D* favor a homogeneous state and which favor a patterned state. We consider the discrete **k** = 2*π*(*n***e**_*x*_ + *m***e**_*y*_)/*L, m, n* = 1, 2, 3,… that satisfy periodic boundary conditions and calculate the phase boundary numerically (Fig. 2A, dashed curve).

**Fig. 2.**
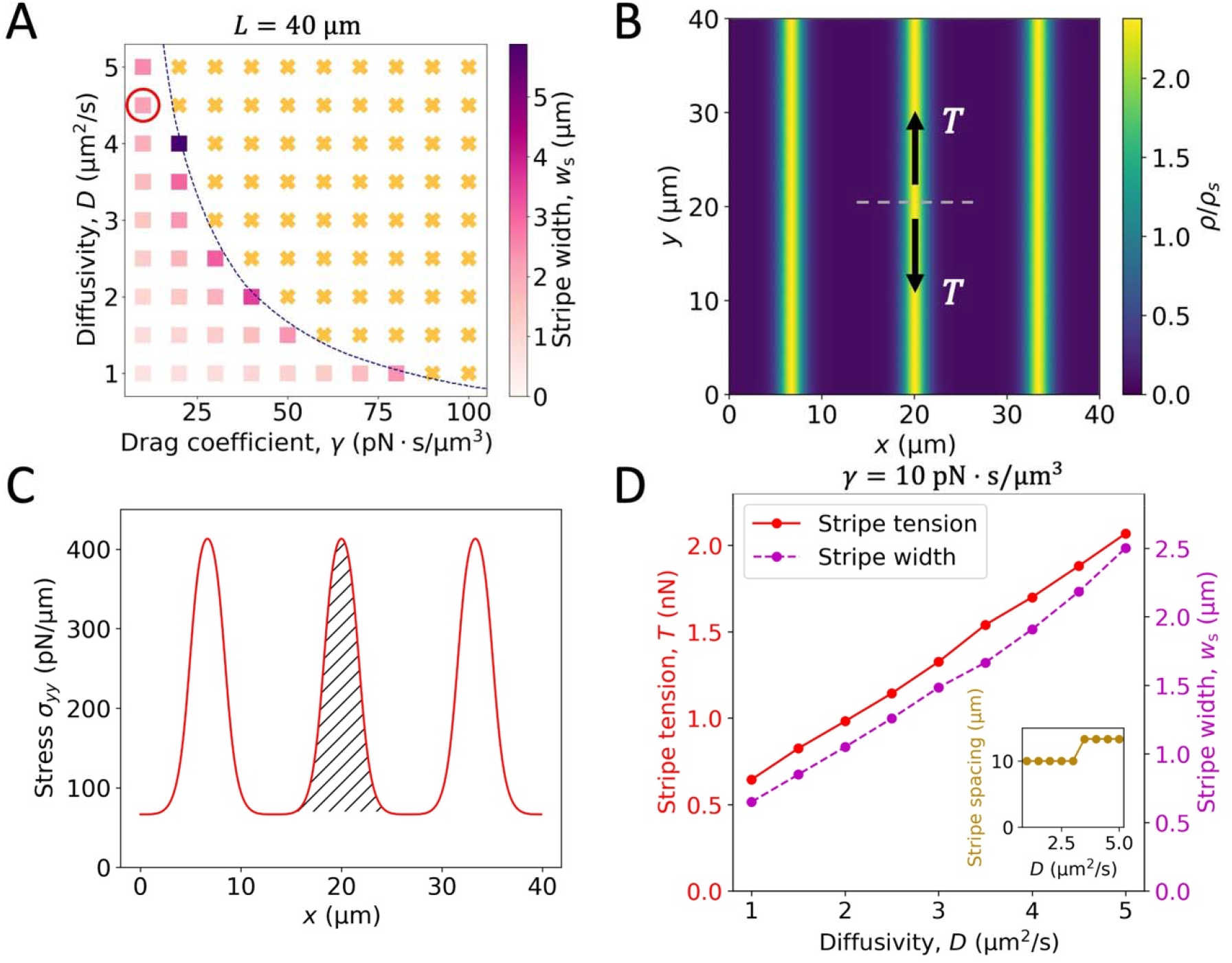
(A) A phase diagram showing stripe width *w_s_* as a function of the drag coefficient *γ* and the actomyosin diffusivity *D. w_s_* is defined as the full width at half maximum (FWHM) of the density profile *ρ*(*x*). Colored squares and crosses are numerical solutions, and the dashed curve represents the phase boundary given by linear stability analysis. Orange crosses, homogeneous steady state; squares, stripe patterns, with stripe width *w*_s_ indicated by the color. The red circle indicates the parameters used in subpanel B. (B) An example of stripe pattern formed on a planar cortex with parameters. The black arrows indicate the tension of the middle stripe. (C) Plot of the stress along the stripe *σ_yy_*(*x*) of the pattern shown in subpanel B. The integrated area (hatched) gives the stripe tension *T*. (D) Plots of stripe tension, stripe width and stripe spacing as a function of the actomyosin diffusivity.

### Striped patterning in the actomyosin cortex

To investigate the patterns that form when the homogeneous state becomes unstable, we solve the full nonlinear equations (1–5) numerically within a flat square with side *L* = 40 μm using the Fourier spectral method. For the initial condition, we apply a perturbation varying only in the *x* direction, a Gaussian of width 0.3 μm and amplitude 0.01*ρ_s_*. The drag coefficient *γ* is scanned from 10 – 100 pN· s/μm^3^ and the diffusivity *D* is scanned from 1 – 5 μm^2^/s.

We find that the cortex spontaneously forms striped patterns when inequality (10) is satisfied (Fig. 2A–B, and S1A–B). Patterns formed by the following mechanism: small perturbations away from the homogeneous state grow because regions with higher density have higher active stress, causing actomyosin material to flow away from regions of low density and accumulate in the higher-density regions. Thus, higher-density regions become even higher in density, in a positive feedback loop.

Fig. 2A shows how the stripe width *w_s_* depends on the diffusivity *D* and drag coefficient *γ*. The stripe tension is defined as the pulling force along the length of the stripe. As *w_s_* increases, there is more actomyosin material in the stripe and thus the stripe exerts higher tension. Fig. 2D shows both the stripe width and the stripe tension grow monotonically with increasing diffusivity *D*, and the stripe spacing only increases very slightly, consistent with the predictions of linear stability analysis, Eq. (8). With higher *D*, density peaks are broader and lower in amplitude. Wider stripes exert higher tension because the decrease in the peak density does not affect the local active stress much since the peak density is significantly greater than the half-saturation density *ρ_s_*. The stripe tension *T* is calculated by integrating the stress *σ_yy_*(*x*) over the region where *σ_yy_*(*x*) is more than 1% above the minimum value (Fig. 2C).

To compare these solutions with those presented by Hannezo et al., we repeated our simulations with increased drag coefficient *γ* and decreased diffusivity *D* used by these authors. With these parameters, we find patterns consisting of thin stripes with very narrow spacing (Fig. S1A–B), similar to the solutions of Hannezo et al.

### Cortex patterning drives shape change

Our results on a plane illustrate spontaneous pattern formation of the actomyosin cortex, but did not address morphological transitions. Here we apply the model on a curved surface to study the shape change generated by contractile actomyosin forces. Here, we consider the simplest curved surface, an axisymmetric cylinder, as a model cell cortex.

The axisymmetric surface is represented by the parameterization **X**(*θ, z*) = (*R*(*z*) cos *θ*, *R*(*z*) sin *θ, z*), where *R*(*z*) represents the local distance of the surface from the *z*-axis, Fig. 1B. We define *α*(*z*) = arccot*R′*(*z*) as the angle between the surface tangent vector **e**_*s*_ and the radial vector **e**_*r*_, Fig. 1B. With this parametrization, the principal curvatures *κ_θ_* and *κ_s_* are given by

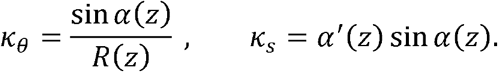

The mean curvature *H* therefore obeys 2*H* = *κ_θ_* + *κ_s_* = sin *α*(*z*) /*R*(*z*) + *α′*(*z*) sin *α*(*z*). The velocity is decomposed into components tangential and normal to the surface, **v** = *v^s^***e**_*s*_ + *v_n_***n**, Fig. 1B.

We first analyze the linear stability of this system and show that spontaneous pattern formation in the cortex drives associated changes in cell shape. Consider an actomyosin cortex anchored to a cylindrical membrane of length *L*, and radius *R*_0_ with periodic boundary conditions and homogeneous steady-state density *ρ*_0_. The pressure in equilibrium is determined by the force balance Eq. (5), 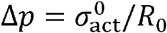, consistent with the Young-Laplace law. This is a steady-state solution because it satisfies both the tangential force balance, Eq. (3) and the normal force balance, Eq. (5). Now we again perturb the density, *ρ* = *ρ*_0_ + *δρ* exp(*ikz* + *λt*). We assume the velocity takes the form *v^s^* = *δv^s^* exp(*ikz* + *λt*) and *v_n_* = *δv_n_* exp(*ikz* + *λt*), and the perturbed shape profile is *R* = *R*_0_ + *δR* exp(*ikz* + *λt*). Expanding the principal curvatures *κ_θ_* and *κ_s_* to first order in *δR* gives 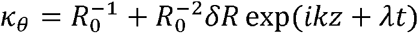 and *κ_s_* = *k*^2^*δR* exp(*ikz* + *λt*).

Following an analogous procedure to that is used in the previous section, we insert the perturbed forms of *ρ, v^s^, v_n_*, and *R* into the constitutive relation Eq. (4) to get the stress *σ^ij^*, and retain first order terms in *δρ, δv^s^, δv_n_* and *δR*. The tangential force balance Eq. (3) and the normal force balance Eq. (5) then yield

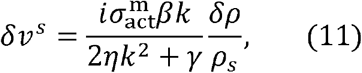

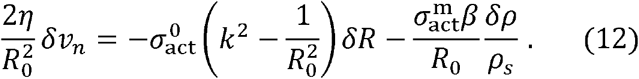

Similarly, retaining only first order terms in *δρ, δv^s^, δv_n_* and *δR*, the density evolution Eq. (1) and the shape dynamics relation Eq. (2) yield

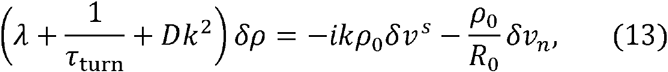

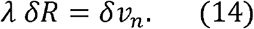

The expressions of Eq. (11–14) show that when density is perturbed, proportional variations in the radius, tangential velocity, and normal velocity are generated. Thus, inhomogeneities in cortical density lead to inhomogeneous flows and shape changes.

Equations (11–14) then give the dispersion relation between *λ* and *k*,

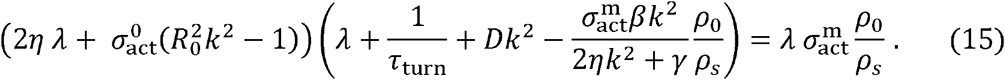

Solving Eq. (15) gives the growth rates *λ* for wavenumbers *k* = 2*nπ*/*L, n* = 1, 2, 3,… that satisfy the periodic boundary condition. The most unstable wavenumber *k*_max_ is that whose growth rate *λ*_max_ has maximal real part.

Using the fact that the homogeneous cylindrically shaped solution is unstable if *λ*_max_ has a non-negative real part leads to the phase diagram of Fig. 3A. This analysis predicts the phase boundary (red curve). For smaller cortex-membrane drag *γ* or smaller actomyosin diffusivity *D*, the cortex forms patterns and furrows the plasma membrane.

**Fig. 3.**
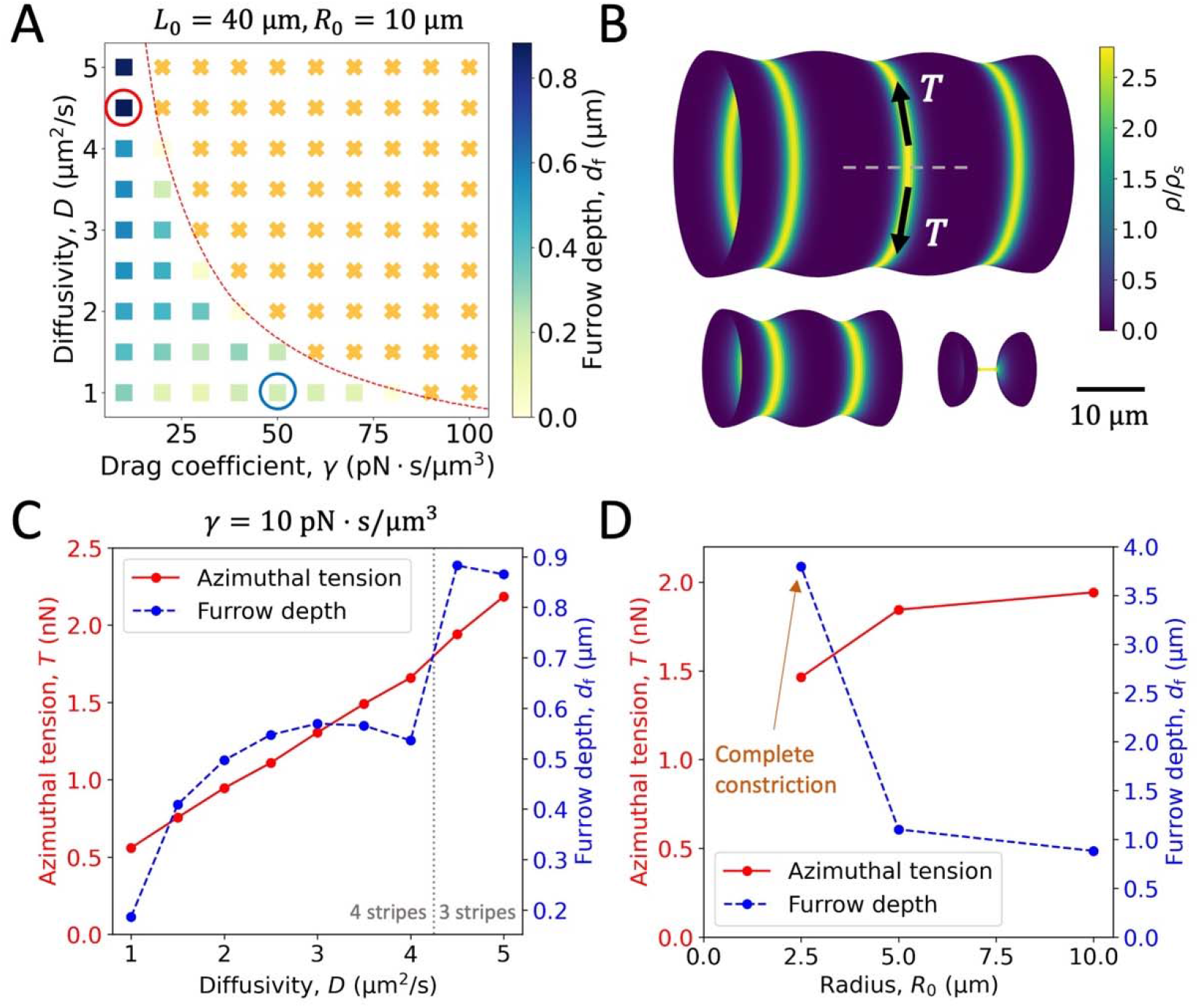
(A) A phase diagram showing furrow depth as a function of the drag coefficient *γ* and the actomyosin diffusivity *D*. The furrow depth *d*_f_ is defined as the mean difference between the peaks and troughs in the shape profile *R*(*z*) Colored squares and crosses are numerical solutions, and the dashed curve represents the phase boundary given by linear stability analysis. The numerical results are consistent with the prediction of the linear stability analysis. Orange crosses, homogeneous steady state and no shape change; colored squares, stripe patterns and furrowed tubule, with furrow depth *d*_f_ indicated by the color. The red circle indicates the parameters used in subpanel B. The blue circle indicates the parameters used in Fig. S2. (B) Examples of patterned cortex that furrows the tubular plasma membrane. The three tubules are of radius *R*_0_ equal to 10 μm, 5 μm and 2.5 μm, respectively, keeping the same aspect ratio *L*/*R*_0_ = 4. The tubule with *R*_0_ = 2.5 μm undergoes complete constriction, and is shown by the last snapshot before constriction. The red arrows indicate the azimuthal tension of the middle stripe. (C) Plots of azimuthal tension and furrow depth with different diffusivity *D*. (D) Plots of azimuthal tension and furrow depth with different radius *R*_0_. In all simulations, parameters in Table 1 unless otherwise indicated.

### Periodic furrowing of the cortex

Next, we want to explain the experimentally observed shape changes on the exocytotic vesicles in *Drosophila* salivary gland cells (Rousso et al., 2016) and the *Drosophila* tracheal tubes with *kkv* mutation (Hannezo et al., 2015). To determine the shape changes triggered by cortical instability, we numerically solved the full nonlinear equations (1–5) on a cylinder with *L* = 40 μm and *R*_0_ = 10 μm for a range of *γ* and *D* values. The initial perturbation is the same Gaussian as for the plane analysis (Fig. 2). Spatial derivatives are approximated using a finite difference scheme based on the discretized arc-length, and the system is evolved in time using the forward Euler method.

The solution showed cortex furrowing, consistent with the predictions of linear stability analysis (Fig. 3A–B, and S2). The cortex aggregates into spatially periodic patterns characterized by ring-like stripes around the axis of the tubular surface separated by depleted regions. These stripes exert larger centripetal contractile forces than the depleted zones, creating furrows on the cylindrical plasma membrane. Thus pattern formation drives shape changes in the cortical surface.

The furrows are deeper for lower drag coefficient *γ* and higher diffusivity *D*. This is reminiscent of the experimental observations of *Drosophila* tracheal tubes, which showed deeper furrowing in *kkv* mutants (Hannezo et al., 2015), despite the tracheal tube is a multi-cellular system. Actin rings in these mutants are thought to have lower drag due to the extracellular matrix being compromised.

We examined the relation between furrow depth and azimuthal tension *T*exerted by a stripe. As diffusivity *D* increases, both azimuthal tension and furrow depth increase, although the increase of furrow depth is not monotonic (Fig. 3C). Higher azimuthal tension of wider stripes generates deeper furrows. The nonmonotonic increase in furrow depth is due to the discontinuous transition from 4 to 3 stripes.

The furrow depth increased when the cylinder was made smaller while fixing the aspect ratio, Fig. 3D. This increase in furrow depth was not driven by an increase in tension since the azimuthal tension decreased only slightly with smaller cylinder radius *R*_0_. At a critical cylinder size (*R*_0_ ~ 2.5 μm), the local radius of the furrow became zero, the furrowed shape became unstable and the cylinder underwent constriction (Fig. 3B, 3D).

## Discussion

Contractile actomyosin systems such as the cell cortex are intrinsically unstable due to their contractile nature: since the local contractile stress is increased in regions of high myosin density, fluctuations in myosin density produce flows toward the regions of highest density, thus increasing the density in a mechanical feedback loop that destabilizes a homogenous distribution of cortical proteins. This effect threatens the continuity of actomyosin structures such as the cytokinetic ring or the cortex and their capacity to produce tension, a threat cells must control. For example, in cytokinetic rings in cell ghosts, the instability is uncontrolled due to lack of component turnover, leading to hierarchical aggregation of myosin II and breakdown of the organization (Thiyagarajan et al., 2021). However, the contractile instability can also be used productively in cells. For example, formation and constriction of the actomyosin contractile ring that divides cells during cytokinesis can be viewed as a regulated contractile instability (Turlier et al., 2014).

The actomyosin cortex develops patterns of variable actin and myosin concentration. Circumferential actomyosin rings in *Drosophila* determine the spatial pattern of folds in tracheal tubes (Hannezo et al., 2015). Actin rings were also implicated in elongation of the *C. elegans* embryo (Priess and Hirsh, 1986). Patterned coats of myosin-II have also been observed on the surface of secretory granules in *Drosophila* salivary glands (Rousso et al., 2016) and mouse exocrine glands (Ebrahim et al., 2019).

Here we asked if these spatial variations in actin and myosin density in the cortex can lead to shape perturbation, since regions of higher myosin density would exert larger contractile forces. Can this lead to morphological transitions from simple shapes to complex shapes with multiple furrows or other features? Indeed, there is experimental evidence that cortical patterns can provoke shape transitions in cells. Shallow folds in the *Drosophila* tracheal tube colocalized with circumferential actin bands (Matusek et al., 2006), and in the *kkv* mutant, these bands created deep furrows (Hannezo et al., 2015). Striped myosin coats on secretory granules in *Drosophila* salivary glands also lead to substantial furrows of the granule during content release (Rousso et al., 2016).

However, whether these stable shape transitions originate in actomyosin contractility has not been demonstrated theoretically. In a recent study of active gel models on dynamically evolving surfaces, bending rigidity was needed to stabilized furrowed geometries (Mietke et al., 2019); however, the effects of bending rigidity are negligible on cellular scales due to the small bending modulus of cell membranes (Dimova, 2014). Here, we demonstrated within an active gel framework that actomyosin contractility can generate stable deformed structures that spatially coincide with the patterns caused by the contractile instability. This may allow cells to use contractile instability for functional advantage.

Pattern formation and, correspondingly, shape change, occurred when diffusivity or drag coefficient were below a threshold determined by other parameters such as cortical contractility, viscosity and turnover time, Eq. (17). The magnitude of the shape change generated by observed patterns, measured by the furrow depth, increased significantly with diffusivity because higher diffusivity gives thicker stripes that exert larger azimuthal tension, and constrict the plasma membrane more. The furrow depth also depended on the initial size of the cylindrical cortex, with deeper furrows in smaller cells.

Our model implemented a finite actomyosin diffusivity in order to stabilize the system against small length scale aggregations that make simulating the active gel model numerically intractable, which is commonly used in other active gel models (Bois et al., 2011b; Hannezo et al., 2015). However, the values used are much bigger than the experimentally measured myosin II diffusivity (about 2 × 10^−5^ μm^2^/s) in the *S. pombe* fission yeast contractile rings (Vavylonis et al., 2008). Diffusivities were also reported to be unmeasurably small in the cell cortex of *C. elegans* early embryo (Nishikawa et al., 2017).

If the diffusive term is unphysical, the question of what prevents runaway aggregation in cells arises. A recent study proposed that it is the rapid myosin turnover that prevents runaway aggregation in fission yeast contractile rings, based on the observed runaway hierarchical aggregation of actomyosin nodes in cell ghosts with no turnover (Thiyagarajan et al., 2021). Another proposed mechanism is that the steric repulsion in high-density regions prevents contraction beyond a maximum density and stabilizes the cortex on the apical surface of *Drosophila* ectoderm cells (Munjal et al., 2015).

## Acknowledgement

This work was supported by National Institute of General Medical Sciences of the National Institutes of Health under award number R01GM086731 to B.O. The content is solely the responsibility of the authors and does not necessarily represent the official views of the National Institutes of Health. The authors have no conflict of interest to declare. We acknowledge computing resources from Columbia University’s Shared Research Computing Facility project.

---

**Fig. S1.**
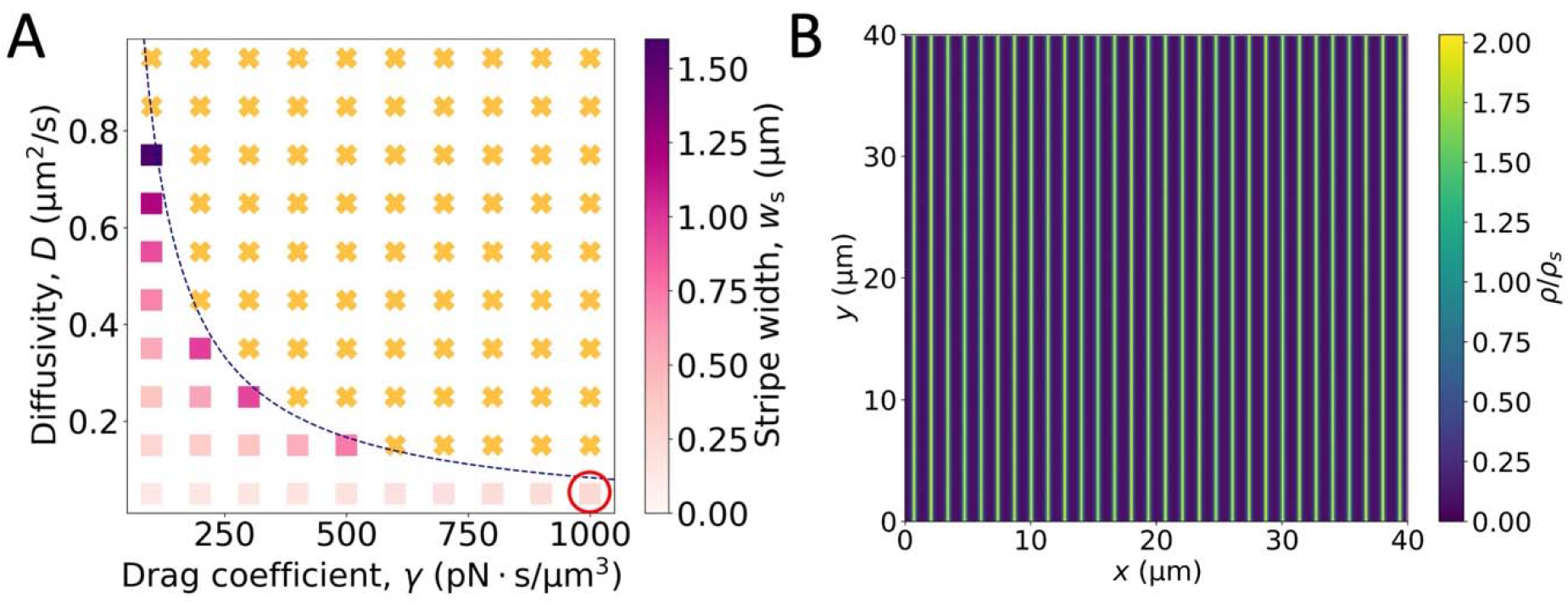
(A) Numerical generated phase diagram with higher drag coefficient *γ* and lower diffusivity *D*. The dashed curve is the phase boundary predicted by the linear stability analysis. The red circle indicates the parameters used in subpanel B. (B) An example of stripe pattern with higher *γ* and lower *D*.

**Fig. S2.**
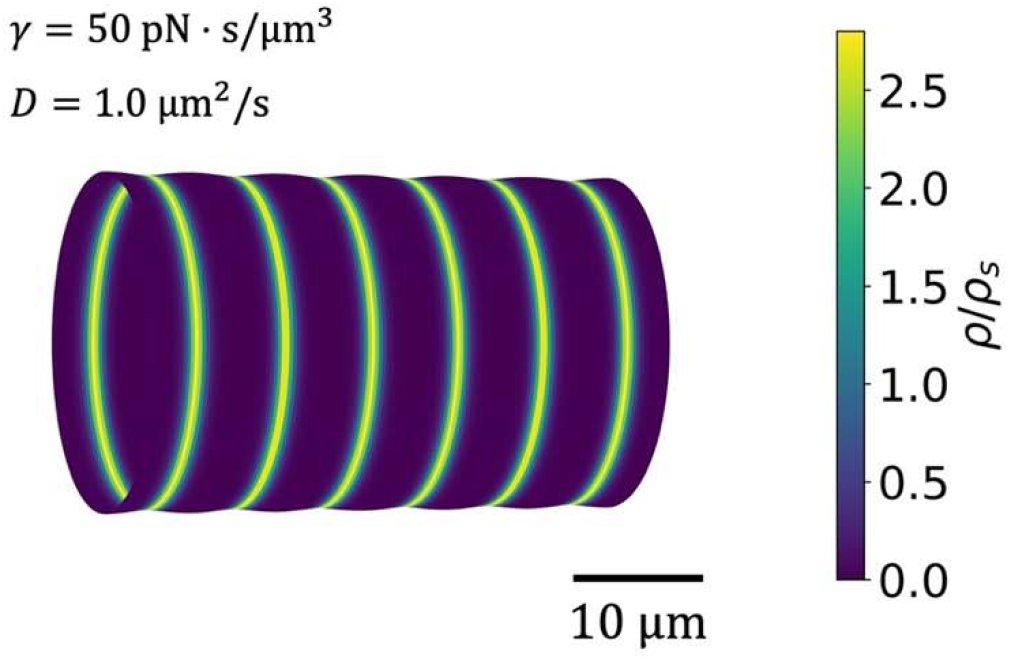
An example of shallow furrows, corresponding to the blue circle in Fig. 3A.

## References

Berthoumieux, H., Maître, J.-L., Heisenberg, C.-P., Paluch, E.K., Jülicher, F., and Salbreux, G. (2014). Active elastic thin shell theory for cellular deformations (New Journal of Physics), pp. 065005.

Bois, J.S., Julicher, F., and Grill, S.W. (2011a). Pattern Formation in Active Fluids. Phys Rev Lett 106.

Bois, J.S., Julicher, F., and Grill, S.W. (2011b). Pattern Formation in Active Fluids. Biophysical Journal 100, 445–445.

Charras, G.T., Yarrow, J.C., Horton, M.A., Mahadevan, L., and Mitchison, T.J. (2005). Non-equilibration of hydrostatic pressure in blebbing cells. Nature 435, 365–369.

Chugh, P., and Paluch, E.K. (2018). The actin cortex at a glance. J Cell Sci 131.

Dimova, R. (2014). Recent developments in the field of bending rigidity measurements on membranes. Adv Colloid Interface Sci 208, 225–234.

Ebrahim, S., Chen, D., Weiss, M., Malec, L., Ng, Y., Rebustini, I., Krystofiak, E., Hu, L., Liu, J., Masedunskas, A., et al. (2019). Dynamic polyhedral actomyosin lattices remodel micron-scale curved membranes during exocytosis in live mice. Nat Cell Biol 21, 933–939.

Evans, E., and Yeung, A. (1989). APPARENT VISCOSITY AND CORTICAL TENSION OF BLOOD GRANULOCYTES DETERMINED BY MICROPIPET ASPIRATION. Biophysical Journal 56, 151–160.

Hannezo, E., Dong, B., Recho, P., Joanny, J.F., and Hayashi, S. (2015). Cortical instability drives periodic supracellular actin pattern formation in epithelial tubes. P Natl Acad Sci USA 112, 8620–8625.

Heer, N.C., Miller, P.W., Chanet, S., Stoop, N., Dunkel, J., and Martin, A.C. (2017). Actomyosin-based tissue folding requires a multicellular myosin gradient. Development 144, 1876–1886.

Maddox, A.S., and Burridge, K. (2003). RhoA is required for cortical retraction and rigidity during mitotic cell rounding. J Cell Biol 160, 255–265.

Matusek, T., Djiane, A., Jankovics, F., Brunner, D., Mlodzik, M., and Mihály, J. (2006). The Drosophila formin DAAM regulates the tracheal cuticle pattern through organizing the actin cytoskeleton. Development 133, 957–966.

Mietke, A., Julicher, F., and Sbalzarini, I.F. (2019). Self-organized shape dynamics of active surfaces. Proc Natl Acad Sci U S A 116, 29–34.

Moeendarbary, E., Valon, L., Fritzsche, M., Harris, A.R., Moulding, D.A., Thrasher, A.J., Stride, E., Mahadevan, L., and Charras, G.T. (2013). The cytoplasm of living cells behaves as a poroelastic material. Nat Mater 12, 253–261.

Moore, T., Wu, S.K., Michael, M., Yap, A.S., Gomez, G.A., and Neufeld, Z. (2014a). Self-organizing actomyosin patterns on the cell cortex at epithelial cell-cell junctions. Biophys J 107, 2652–2661.

Moore, T., Wu, S.K., Michael, M., Yap, A.S., Gomez, G.A., and Neufeld, Z. (2014b). Self-Organizing Actomyosin Patterns on the Cell Cortex at Epithelial Cell-Cell Junctions. Biophysical Journal 107, 2652–2661.

Munjal, A., Philippe, J.M., Munro, E., and Lecuit, T. (2015). A self-organized biomechanical network drives shape changes during tissue morphogenesis. Nature 524, 351-+.

Munro, E., Nance, J., and Priess, J.R. (2004). Cortical flows powered by asymmetrical contraction transport PAR proteins to establish and maintain anterior-posterior polarity in the early C-elegans embryo. Dev Cell 7, 413–424.

Nishikawa, M., Naganathan, S.R., Jülicher, F., and Grill, S.W. (2017). Controlling contractile instabilities in the actomyosin cortex. Elife 6.

Nishimura, T., Honda, H., and Takeichi, M. (2012). Planar Cell Polarity Links Axes of Spatial Dynamics in Neural-Tube Closure. Cell 149, 1084–1097.

Pollard, T.D., and O’Shaughnessy, B. (2019). Molecular Mechanism of Cytokinesis. Annu Rev Biochem 88, 661–689.

Priess, J.R., and Hirsh, D.I. (1986). Caenorhabditis elegans morphogenesis: the role of the cytoskeleton in elongation of the embryo. Dev Biol 117, 156–173.

Rauzi, M., Lenne, P.F., and Lecuit, T. (2010). Planar polarized actomyosin contractile flows control epithelial junction remodelling. Nature 468, 1110–U1515.

Rousso, T., Schejter, E.D., and Shilo, B.Z. (2016). Orchestrated content release from Drosophila glue-protein vesicles by a contractile actomyosin network. Nat Cell Biol 18, 181–190.

Salbreux, G., and Julicher, F. (2017). Mechanics of active surfaces. Phys Rev E 96, 032404.

Thiyagarajan, S., Wang, S., Chew, T.G., Huang, J., Balasubramanian, M.K., and O’Shaughnessy, B. (2021). Myosin turnover controls actomyosin contractile instability. bioRxiv, 2021.2003.2018.436017.

Tinevez, J.Y., Schulze, U., Salbreux, G., Roensch, J., Joanny, J.F., and Paluch, E. (2009). Role of cortical tension in bleb growth. Proceedings of the National Academy of Sciences of the United States of America 106, 18581–18586.

Turlier, H., Audoly, B., Prost, J., and Joanny, J.F. (2014). Furrow Constriction in Animal Cell Cytokinesis. Biophysical Journal 106, 114–123.

Vavylonis, D., Wu, J.Q., Hao, S., O’Shaughnessy, B., and Pollard, T.D. (2008). Assembly Mechanism of the Contractile Ring for Cytokinesis by Fission Yeast. Science 319, 97–100.

